# Genome sequencing links persistent outbreak of legionellosis in Sydney to an emerging clone of *Legionella pneumophila* ST211

**DOI:** 10.1101/144790

**Authors:** VJ Timms, R Rockett, NL Bachmann, E Martinez, Q Wang, SC-A Chen, N Jeoffreys, PJ Howard, A Smith, S Adamson, R Gilmour, V Sheppeard, V Sintchenko

## Abstract

The city of Sydney, Australia, experienced a persistent outbreak of *Legionella pneumophila* serogroup 1 (Lp1) pneumonia in 2016. To elucidate the source and bring the outbreak to a close we examined the genomes of clinical and environmental Lp1 isolates recovered over 7 weeks. A total of 48 isolates from patients and cooling towers were sequenced and compared using SNP-based, core-genome MLST and pangenome approaches. All three methods confirmed phylogenetic relatedness between isolates associated with outbreaks in the Central Business District (March and May) and Suburb 1. These isolates were designated “Main cluster” and consisted of isolates from two patients from the CBD March outbreak, one patient and one tower isolate from Suburb 1 and isolates from two cooling towers and three patients from the CDB May outbreak. All main cluster isolates were sequence type ST211 which has only ever been reported in Canada. Significantly, pangenome analysis identified mobile genetic elements containing a unique T4ASS that was specific to the main cluster and co-circulating clinical strains, suggesting a potential mechanism for increased fitness and persistence of the outbreak clone. Genome sequencing was key in deciphering the environmental sources of infection among the spatially and temporally coinciding cases of legionellosis in this highly populated urban setting. Further, the discovery of a unique T4ASS emphasises the potential contribution of genome recombination in the emergence of successful Lp1 clones.

## Introduction

Between February to May 2016, Sydney, Australia experienced several apparent outbreaks of human legionellosis. The particular features of these outbreaks were that they spanned three precincts within the Greater Sydney region, 18 km apart over seven and a half weeks and included the Central Business District (CBD), an area visited by over 610,000 visitors per day. The agent responsible was found to be *Legionella pneumophila* serogroup 1 (Lp1), a ubiquitous organism with a natural habitat of water reservoirs and amoebae that also sporadically infects humans and during this period it was responsible for the death of two patients.

The genus *Legionella* contains over 60 species. All species are found in the environment and about half of these can cause a life-threatening illness in humans. *L. pneumophila* is the major pathogen of this group and is further subdivided into 16 serogroups with the majority (80-84%) of legionellosis cases caused by Lp1 (Ambrose et al., 2014)(Kozak-Muiznieks et al., 2014)(Raphael et al., 2016). Since the first description of Legionnaires disease in 1976 when a large outbreak of pneumonia occurred at an American Legion conference in Philadelphia (Fraser et al., 1977), this infection has been implicated in multiple sporadic cases and community outbreaks globally, most commonly in males over 50, people with co-morbidities and smokers (Frieden, Jaffe, Stephens, & Thacker, 2011).

The *Legionella* are highly recombinant bacteria with the latest recombination/mutation rate reported at 47.93 per site (Sánchez-Busó, Comas, Jorques, & González-Candelas, 2014)(Gomez-Valero et al., 2011) coupled with a suite of mobile genetic elements, including plasmids and pathogenicity islands (Khodr et al., 2016; Wee, Woolfit, Beatson, & Petty, 2013)(Mercante, Morrison, Desai, Raphael, & Winchell, 2016)(Chien et al., 2004). These mobile genetic elements encode virulence and fitness factors, such as type IV secretion systems, known to be imperative for intracellular replication and survival in both amoebae and human macrophages (Gomez-Valero & Buchrieser, 2013)(Voth, Broederdorf, & Graham, 2012). While gene sequencing based typing (SBT) methods have been used for outbreak investigation (Gaia et al., 2005), the limited resolution of SBT have led to a wider application of high resolution whole genome sequencing (WGS) in the investigation of community outbreaks of legionellosis. WGS technological advancements now provide increased discrimination of outbreak isolates especially for endemic clones such as ST1 (Bartley et al., 2016; Reuter et al., 2013)(Graham, Doyle, & Jennison, 2014). This study was aimed to elucidate the source and probable increased persistence of the complex Sydney 2016 outbreak by examining the genomes of clinical and environmental *L. pneumophila* isolates collected over that peroid.

## Methods

### Bacterial isolates and serogrouping

Between February and May 2016, cases were identified by positive urinary Lp1 antigen results and/or positive respiratory tract cultures for Lp1 and clinical findings (details available on request). Clinical and environmental Lp1 isolates from three geographically distinct locations, the Central Business District (CBD), 18 km south of the CBD (Suburb 1) and 12 km west of the CBD (Suburb 2), were studied (Table 1 and Figure 1). Isolates from CBD spanned February to March 2016 (CBD March outbreak) and from April to May (CBD May outbreak). There was a period of six weeks where no cases from the CBD were detected between March and April. Environmental isolates from cooling towers were referred from the Forensic and Analytical Science Service, whilst clinical isolates were cultured from sputum samples or bronchoalveolar lavage fluid referred to the Centre for Infectious Diseases and Microbiology Laboratory Services, ICPMR-Pathology West, Westmead Hospital, Sydney. Isolates were grown on buffered charcoal yeast extract (BCYE) agar and incubated at 37°C for up to 7 days and identified by standard phenotypic characteristics (Versalovic et al., 2011). Serogrouping of each isolate was performed using the Legionella Latex Test (Oxoid, ThermoFisher Scientific, North Ryde, NSW, Australia). The full co-ordinated public health response is detailed in the Public Health Investigation into the Legionella outbreaks in Sydney CBD (Griffiths et al., 2016).

**Figure 1:**
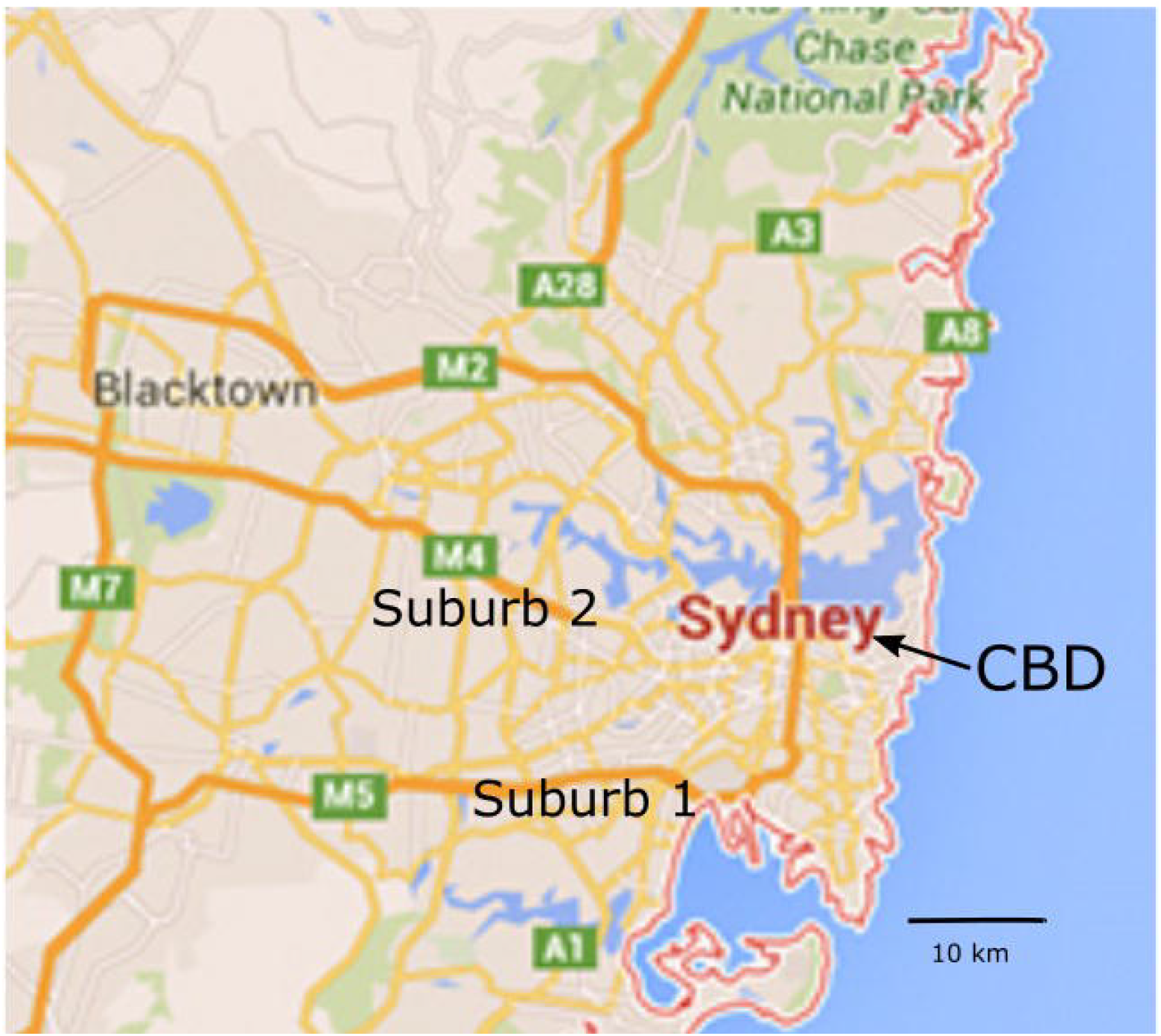
Map of metropolitan Sydney indicating locations of the outbreak.

**Table 1.** *Legionella pneumophila* serogroup 1 isolates included in the study.

| **Isolate identification number** | **Origin** | **Case number in outbreak** | **Association with location** | **SBT type** |
| --- | --- | --- | --- | --- |
| BC7 | Clinical | Case 1 | CBD March* | 211 |
| BC10 | Clinical | Case 2 | CBD March | 42 |
| BC5 | Clinical | Case 3 | CBD March | 211 |
| Reference control 1 | Clinical | ATCC33152 |  |  |
| Reference control 2 | Clinical | ATCC33215 |  |  |
| BC2 | Clinical | Case 1 | CBD March | 211 |
| BC1 | Clinical | Case 1 | CBD March | 211 |
| BC9 | Clinical | Case 2 | CBD March | 42 |
| Reference 2013 | Clinical | Reference case 1 | Not related | 762 |
| Reference 2015 | Clinical | Reference case 2 | Not related | ND |
| BE3 | Environmental | Tower 1 | CBD March | ND |
| BE11 | Environmental | Tower 2 | CBD March | 1 |
| BE12 | Environmental | Tower 3 | CBD March | 1 |
| BE13 | Environmental | Tower 4 | CBD March | 1 |
| BE7 | Environmental | Tower 5 | CBD March | 1 |
| BE5 | Environmental | Tower 6 | CBD March | 1 |
| BE17 | Environmental | Tower 6 | CBD March | 1 |
| BE20 | Environmental | Tower 7 | CBD March | 284 |
| BE19 | Environmental | Tower 8 | CBD March | 1 |
| BE14 | Environmental | Tower 9 | CBD March | 1 |
| BE10 | Environmental | Tower 9 | CBD March | 1 |
| BE6 | Environmental | Tower 9 | CBD March | 1 |
| BE8 | Environmental | Tower 10 | CBD March | 1 |
| RC3 | Clinical | Case 4 | Suburb 1 | ND |
| RC1 | Clinical | Case 5 | Suburb 1 | 211 |
| RC2 | Clinical | Case 5 | Suburb 1 | 211 |
| RE6 | Environmental | Tower 11 | Suburb 1 | 211 |
| RE4 | Environmental | Tower 11 | Suburb 1 | 211 |
| RE7 | Environmental | Tower 11 | Suburb 1 | 211 |
| RE5 | Environmental | Tower 11 | Suburb 1 | 211 |
| RE18 | Environmental | Tower 12 | Suburb 1 | 1 |
| RE2 | Environmental | Tower 12 | Suburb 1 | 1 |
| RE16 | Environmental | Tower 12 | Suburb 1 | 1 |
| RE15 | Environmental | Tower 12 | Suburb 1 | 1 |
| RE3 | Environmental | Tower 12 | Suburb 1 | 1 |
| RE1 | Environmental | Tower 10 | Suburb 1 | 284 |
| BC6 | Clinical | Case 6 | CBD May | 211 |
| BC4 | Clinical | Case 7 | CBD May | 211 |
| BC3 | Clinical | Case 7 | CBD May | 211 |
| BC8 | Clinical | Case 8 | CBD May | 211 |
| BE4 | Environmental | Tower 13 | CBD May | ND |
| BE1 | Environmental | Tower 14 | CBD May | 211 |
| BE9 | Environmental | Tower 9 | CBD May | 1 |
| BE2 | Environmental | Tower 14 | CBD May | 211 |
| GC3 | Clinical | Case 9 | Suburb 2 | 1983 |
| GC2 | Clinical | Case 10 | Suburb 2 | ND |
| GC1 | Clinical | Case 11 | Suburb 2 | 84 |
| GC4 | Clinical | Case 12 | Suburb 2/CBD May | ND |

\*Abbreviations: CBD March: Isolates from the Central Business District during February to
March 2016. CBD May, isolates from the Central Business District between April to May
\*\*ND – Not Determined due to one or more alleles being new or combination of alleles being
new.

For whole genome analysis, the type strains *L. pneumophila* ATCC 33152 (Philadelphia AE017354) and *L. pneumophila* ATCC33215 (Chicago 2 933085) were included for comparison of assembly and typing pipelines. In addition, two Lp1 clinical isolates from previous unrelated outbreaks in Sydney (2013 Reference and 2015 Reference) were included as unrelated outgroups.

### DNA extraction and WGS

Genomic DNA was extracted from pure cultures using the DNeasy Blood & Tissue Kit (QIAGEN, Chadstone, VIC, Australia) with a 3 hour Proteinase K digestion at 56°C. The quality of each DNA sample was measured using a Nanodrop ND-1000 spectrophotometer (Nanodrop Technologies, ThermoFisher Scientific) and the ratios 260/280 nm and 260/230 nm inspected for the presence of organic matter and solvent residues, respectively. DNA purity was determined as satisfactory if the 260/280 ratio fell between 1.6 and 2.2 and the 260/230 ratio fell between 1.8 and 2.2. Purity of each DNA sample was also inspected on 1.5% agarose gel. Paired-end indexed libraries of 150 bp in length were prepared from an input of 1 ng of purified DNA with the Nextera XT library preparation kit (Illumina, Scoresby, VIC, Australia) as per manufacturer’s instructions. DNA libraries were then sequenced using the NextSeq 500 (Illumina).

### Genome assembly and analysis

To identify SNPs from whole genome sequencing data, FASTQ files were imported into Geneious (version 8.0.4) and mapped to a curated reference of *L. pneumophila* Philadelphia (accession number NC_002942) using the bwa plugin (version 0.7.10) with bacteriophage, insertion sequences and other repeat regions removed according to a previous publication (Coil et al., 2008). Quality based variant detection was performed using CLC Genomics Workbench v 7.0 (CLC bio Aarhus, Denmark). Variant detection thresholds were set for a minimum coverage of 10 and minimum variant frequency of 75%. SNPs were excluded if they were in regions with a minimum fold coverage of <10, within 10-bp of another SNP or <15-bp from the end of a contig. Maximum likelihood phylogenetic trees were constructed from SNP matrices using the GTR model with 100 bootstrap replications.

Sequencing reads were assembled with Spades (Bankevich et al., 2012) and annotated with Prokka (Seemann, 2014). Sequence-based typing (SBT) was performed with seven loci by uploading identified alleles to the database and obtaining loci number (Gaia et al., 2005). In addition, core genome multi-locus sequence typing (cgMLST) was conducted using the Ridom SeqSphere software (Qiagen) employing gene definitions determined from closed or complete *L. pneumophila* genomes from NCBI GenBank (Moran-Gilad et al., 2015). Using these criteria, the core genome was determined to be 1530 genes with an accessory genome of 1370 genes.

Further pangenome assessment and visualisation was performed using the Roary pipeline (Page et al., 2015) that includes alignment using MAFFT (Katoh, Misawa, Kuma, & Miyata, 2002) and tree building with FastTree (Price et al., 2010). Specific protein and nucleotide comparisons were made (including the unique T4SS) to all available Legionella genomes on NCBI using the relevant Blast database (https://blast.ncbi.nlm.nih.gov/Blast.cgi).

The genomic data have been deposited in the NCBI Sequence Read Archive (SRA) (http://www.ncbi.nlm.nih.gov/Traces/sra/) under accession number (TBA).

## Results

Four outbreaks of severe community-acquired pneumonia in adults with positive urinary antigen for Lp1 were identified in metropolitan Sydney in the first half of 2016. Nine human cases were linked to the CBD March outbreak and six cases in the CBD May outbreak, all with common exposure links to the Sydney CBD. Detailed epidemiological information on the CBD cases can be found in the Public Health Investigation Report (Griffiths et al., 2016). Four cases were linked in April to Suburb 1. Also in May, five cases were linked to Suburb 2. One case (Case 12) in May was linked to both Suburb 2 and the CBD (Figures 2 and 3). Lp1 was cultured from respiratory samples of twelve patients and from fourteen cooling towers. All clinical and environmental isolates collected, together with two control strains and two Lp1 isolates obtained from epidemiologically unrelated cases were subjected to genome sequencing (Table 1). All 48 genomes were *de novo* assembled and their genome size ranged between 3.2-3.4 Mb with a G+C of 39%. The core genome as determined by the Roary pipeline, of this dataset consisted of 2181 genes. The accessory genome had 2273 genes out of a total of 5919 genes (including 99 soft core genes and 1366 shell genes that were not counted in the core and accessory respectively).

**Figure 2:**
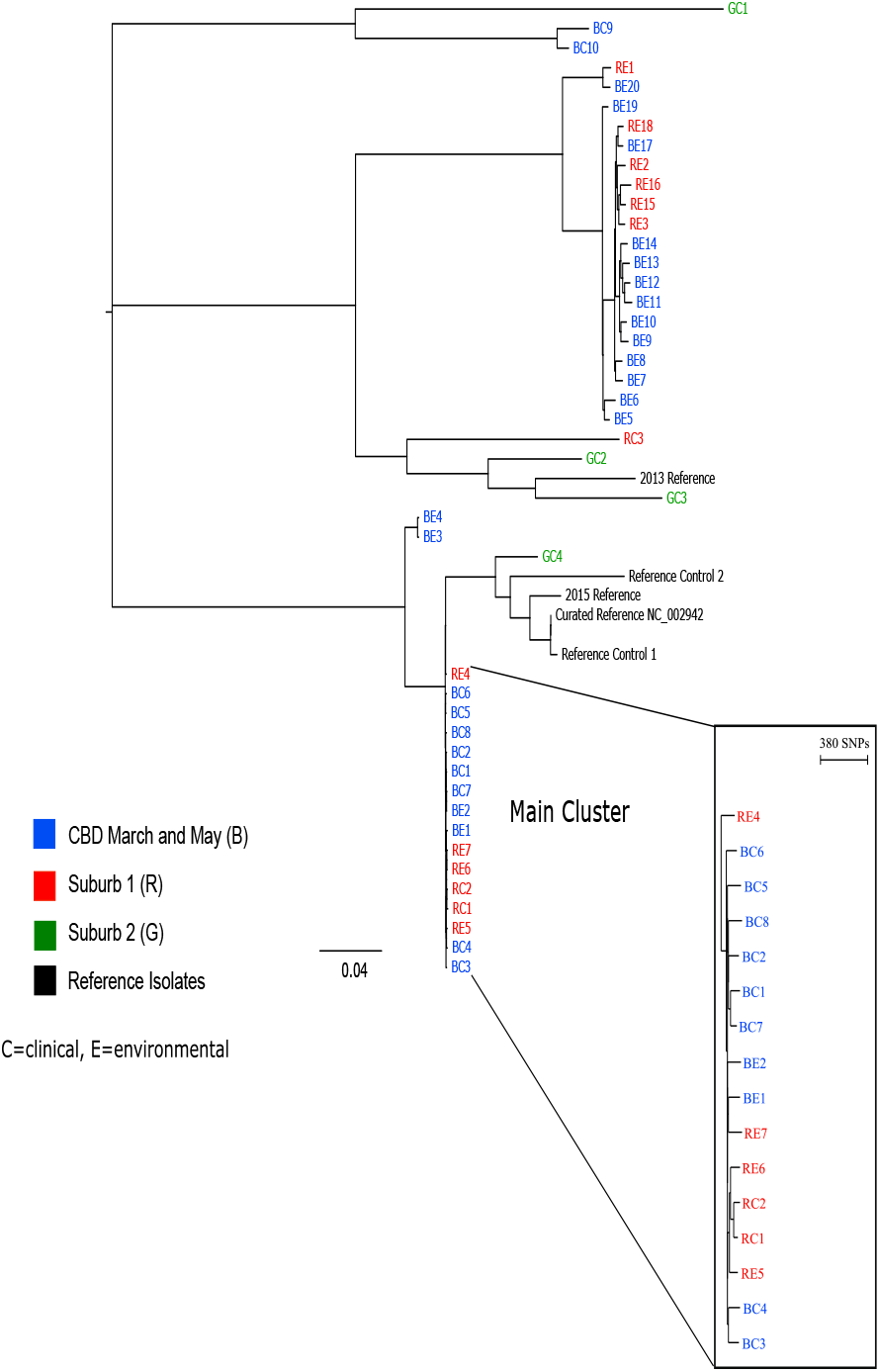
SNP-based mapping phylogeny of all outbreak isolates between February-May 2016 in Sydney.

**Figure 3:**
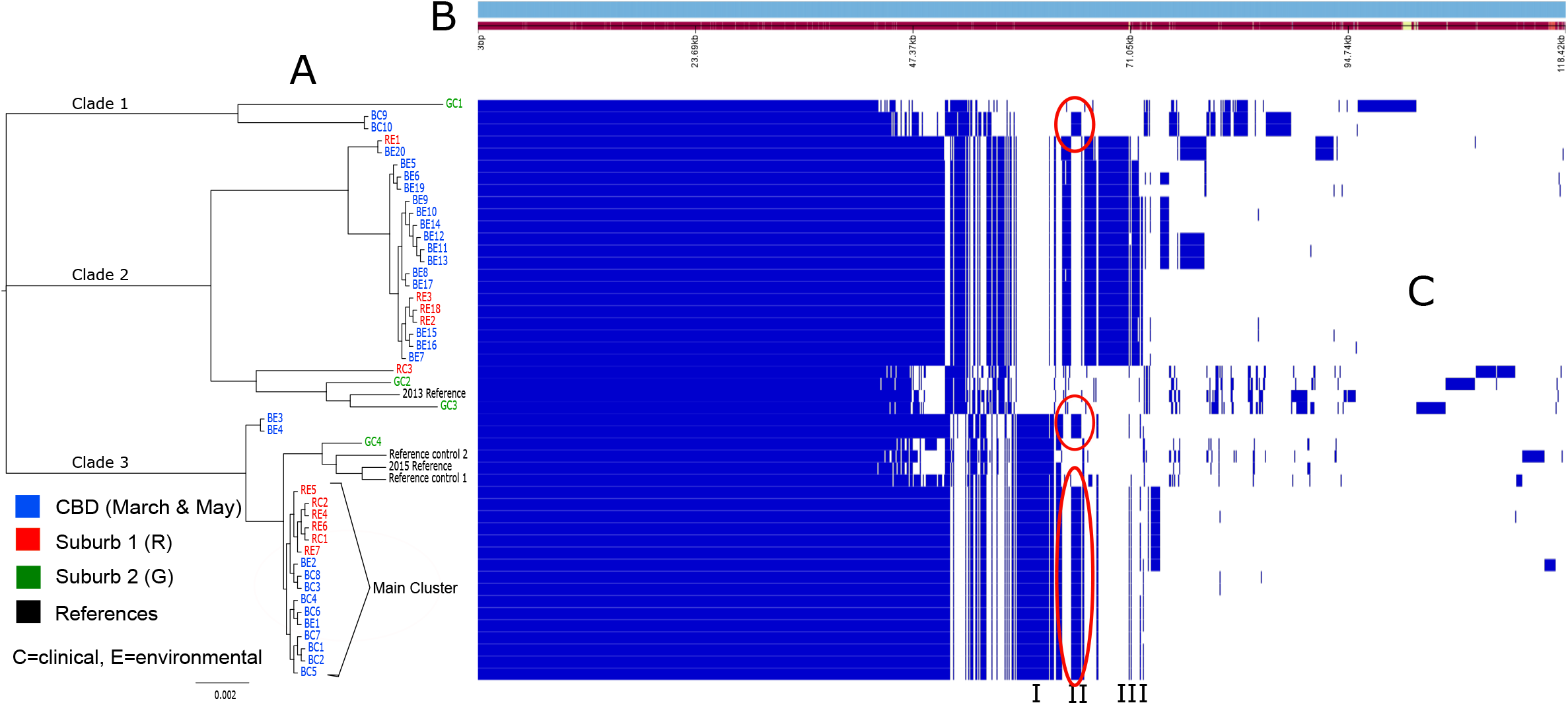
The pan genome of the complete Sydney 2016 dataset (n=48). **A**. Maximum likelihood tree showing outbreak cluster as clade 3. **B**. Pangenome sorted from core genes on the left to accessory genes to the right. **C**. Heatmap showing presence (blue) and absence (white) of genomic regions. The T4SS unique to the outbreak cluster (Region II), the outlier (BC9 and BC10 in blue) and related environmental strains (BE3 and BE4 in blue) are indicated by red circles. Region I demonstrates genes that were unique to clade 3 including reference isolates and the outbreak cluster. Region III indicates genes that were unique to the outbreak cluster and related environmental strains BE3 and BE4 only.

Genome comparisons based on SNP-based mapping, cgMLST and pangenome produced congruent results and confirmed phylogenetic relatedness between isolates associated with outbreaks in the CBD (March and May) and Suburb 1 (Figures 2 and 3 and Supplementary Figure 1) and these isolates were designated “Main cluster”. The CBD March outbreak showed two patients, Cases 1 and 3 clustered together (three isolates were recovered from one patient) (Figures 2 and 3). For the next outbreak in Suburb 1, one patient clustered with one tower and in the CDB May outbreak, isolates from two cooling towers and three patients were of the same genetic cluster. All isolates from the main cluster were also typed as ST211. Isolates from all patients linked to Suburb 2 (Cases 9, 10, 11 and 12) and from one patient linked to each of the CBD (Case 2) and Suburb 1 outbreaks (Cases 4) appeared to be genomically distinct and not related to the main cluster (Figure 2). The maximum SNP difference between outbreak isolates was 87 between CBD tower isolates and Suburb 1 clinical isolates. There were 44 SNP differences between Suburb 1 tower isolates and Suburb 1 clinical isolates. Amongst all clinical isolates from the outbreak cluster, there was an average difference of 66 SNPs and isolates from the same case had less than 45 SNPs. The two towers harbouring ST211 strains were approximately 800m apart on different buildings.

Pangenome analysis further revealed three regions of interest related to the outbreak cluster (Figure 3). The first (Region I) was a large region of 178 genes that was present in the bottom clade only and included the main cluster. Region I was also found in all reference strains, the two past outbreak isolates, closely related environmental strains BE3 and BE4 and an outlier GC4. Region I included the genes for Dot/Icm, *luxR* Ankyrin repeats and beta-lactamase among others indicating that this was a Type IV B secretion system (T4BSS). The second region (Region II) consisted of 58 genes that were unique to the outbreak cluster, the outlier patient isolates BC9 and BC10 co-circulating at the same time and two closely related environmental isolates, BE3 and BE4 (Figure 3). This was found to carry another T4SS, an A F-type secretion system (T4ASS) (Supplementary Table 1) and this region had a higher G+C content of 42% compared to the rest of the genome. Using BLAST this region was found to have between 99-100% homology to similar conjugative elements in only four other *L. pneumophila* Lp1 strains, two closed genomes, LPE509 (NC_020521.1) and C1-S (CP05932.1) and two further clinical isolates, one ST37 isolated in 2003 in the U.K. (LT632617.1) and the other an ST42 clinical isolate from Germany (LT632616.1). This unique region is 73-kb with an insertion site adjacent to three tRNA genes (tRNA^Arg^, tRNA^Lys^, tRNA^Lys^). Two more small regions were unique to the outbreak cluster and closely related isolates BE3 and BE4. These regions denoted as Region III contained mainly hypothetical proteins, acytransferase genes and a calcium transporting ATPase.

## Discussion

Our findings indicated that cases amongst four temporally and spatially separate outbreaks of legionellosis that occurred in quick succession in metropolitan Sydney in 2016 were caused by a common clone of Lp1 ST211. The geographical distance between sites of potential exposure was much larger than previously recognised (Knox et al, 2016) and this coupled with the temporal differences between these outbreaks led to the initial assumption that they were related to separate breaches of environmental health controls. However, phylogenetic analysis on both clinical and environmental isolates from these outbreaks by three independent genomic approaches confirmed that a common Lp1 clone was responsible across three of the four outbreaks. The CBD March outbreak was consonant with the Suburb 1 outbreak 18 km away and both of these were genetically related to the CBD May outbreak that occurred six weeks later. Factors such as cooling tower maintenance and other environmental considerations may have contributed to this wave of Lp1 disease in Sydney.

This report describes the direct comparison of the resolution power of SNP-based and core genome combined with a pangenome based analyses in the investigation of Lp1 outbreaks (David, Mentasti, et al., 2016). Our experience suggested that these three approaches have comparable discrimination power and should be supplemented by the analysis of mobile genetic elements given the high recombination rates in Lp1 (Wang et al., 2015)(Sánchez-Busó et al., 2014). Importantly, the outbreaks in Sydney demonstrated the presence of Lp1 ST211 in the Southern Hemisphere. This ST has not been reported in the USA or Europe and was thought to reside exclusively in the colder climate of Canada where it was first identified in 1989 (Tijet et al., 2010). In Ontario, ST211 was initially thought to be contributing to sporadic cases which peaked in 1999, however, further investigation found that it was responsible for 12.5% of isolates, including eight times in the same hospital. ST211 has since become one of the most persistent and predominant STs in Ontario and along with other STs, has almost completely replaced the previously dominant Lp1 ST1. There is also a suggestion that ST1 may be the predominant ST in Australia (Graham et al., 2014) and further investigations will determine if ST211 is replacing ST1 as the dominant clone in Sydney.

In addition to the assessment of potential relationships between clinical cases and environmental sources, WGS of *Legionella* isolates enables the examination of diversity and genomic structures that may have augmented their persistence in the environment (Sánchez-Busó et al., 2016). Our pangenome analysis revealed a unique conjugative Tra (F-type) element carrying a T4ASS and this was present in all outbreak isolates but more significantly it was also found in closely related environmental isolates and isolates that infected Case 2 at the same time as the CBD March outbreak. Interestingly, this element had 99% nucleotide homology to four previously reported clinical strains from the U.S.A. and Europe and an environmental isolate from Japan, although none of these other isolates are ST211 but are STs in the top five of outbreak causing clones (David, Rusniok, et al., 2016).

The T4SS of *Legionella* are part of the dynamic accessory genome, known to contribute to fitness and virulence and play a crucial role in intracellular replication and survival (Khodr et al., 2016)(Rolando & Buchrieser, 2014; Schroeder et al., 2010; Voth et al., 2012). Unique T4ASS have been found in previous outbreak isolates (Graham et al., 2014) including from a recent outbreak in Western Canada, unique in its dry, cold conditions that were initially thought to be too harsh for survival of *L. pneumophila* (Knox et al., 2016). The element described in this study contained genes homologous to the Lvh region of other *L. pneumophila* strains and genes from this region are thought to assist in intracellular replication (Bandyopadhyay, Liu, Gabbai, Venitelli, & Steinman, 2007)(Bandyopadhyay, Lang, Rasaputra, & Steinman, 2013). In addition, it also contained *csrA*, a carbon storage regulator known to control the switch from replicative to transmissive phase in *Legionella* which is crucial for efficient replication both *in vitro* and intracellularly (Fettes, Forsbach-Birk, Lynch, & Marre, 2001)(Forsbach-Birk, Mcnealy, Shi, Lynch, & Marre, 2004; Molofsky & Swanson, 2003). It is possible that these genomic features reflect high recombination potential and fitness of Lp1 ST211 and help to explain its emergence as an outbreak clone. Earlier reports attempted to define outbreak strains from non-outbreak strains and genome sequencing has identified recombination as a major contributor to Lp1 variability which occurs across all strains, although some STs appear to have different recombination rates (Sánchez-Busó et al., 2014). The genomes described in this study are the only publicly available ST211 genomes that can be used for further outbreak comparisons given the isolates from Canada were typed using the SBT scheme only and not with WGS.

The identification of elements such as the T4SSA described here can alert investigators to the presence of a *L. pneumophila* clone with possible superior virulence and persistence abilities. In addition, given the dependency of *L. pneumophila* on T4SS for intracellular survival and replication, these systems offer an potential target for antibacterial agents and vaccines (Voth et al., 2012).

In conclusion, SNP-based, core genome and pan genome based analyses of Lp1 isolates can assist in deciphering and confirming transmission pathways during the investigation of complex outbreaks of legionellosis. Comparative genomics of clinical and environmental isolates of Lp1 suggested that the emerging ST211 clone was responsible for three sequential outbreaks of legionellosis in metropolitan Sydney separated geographically and temporally over seven weeks. Significantly, pangenome analysis identified mobile genetic elements that included a T4ASS in the outbreak strain and co-circulating strains that may have augmented the fitness and persistence of the outbreak clone in the environment.

## Acknowledgements

The authors thank public health professionals from NSW Health and the City of Sydney, and laboratory scientists from NSW Health Pathology for providing clinical and environmental isolates and their assistance in collecting epidemiological and environmental information.

---

**Supplementary Table 1.** Genes present in T4SS found exclusively in outbreak cluster, BC9, BC10 (Case 2), BE3 and BE4.

**Supplementary Figure 1:** cgMLST phylogeny of all outbreak isolates between February-May 2016 in Sydney.

